# Survival of NASA-cleanroom microbial isolates under simulated space and Martian conditions

**DOI:** 10.1101/2025.10.15.682532

**Authors:** Atul M. Chander, David J. Burr, Severin Wipf, Ruben Nitsche, Gretchen Fujimura, Wayne Schubert, Nitin K. Singh, Justin J. Bell, Alexander Brandl, Michael M. Weil, Andreas Elsaesser, Kasthuri Venkateswaran

## Abstract

Planetary protection hinges on understanding microbial survival following sterilization procedures, the stressors of space travel, and exposure to extraterrestrial environmental conditions. This study identified 23 fungal strains isolated from NASA-spacecraft assembly cleanrooms, capable of surviving ultraviolet radiation exposure. Using experimental simulation facilities, we conducted a comprehensive assessment of microbial survivability and morphology on the most resilient spacecraft-associated microorganisms.

*Aspergillus calidoustus* demonstrated remarkable survival under simulated Martian conditions, withstanding up to 1440 minutes of Martian solar irradiation, Mars atmospheric pressure and composition, and the presence of Martian regolith. Lethality only occurred under combined irradiation and cooling to -60°C (the mean Mars surface temperature), emphasising the synergistic effect of these conditions. Furthermore, *A. calidoustus* survived long-duration neutron radiation exposure (replicating ionizing space radiation doses) and dry-heat sterilization (typically used for spacecraft components).

This is the first study to perform an end-to-end evaluation of eukaryotic microbial survival across conditions that occur during preparation for, travel to, and robotic exploration of Mars. The experimental facilities and chronic exposure methods utilized offer a biologically meaningful model for understanding microbial risks during long-duration space missions. The capacity for fungal conidia to survive multiple space-relevant conditions suggests their potential as forward contaminants, capable of being transported to and persisting on Mars. As current spacecraft sterilization protocols prioritize bacterial spores, this research highlights a critical gap in planetary protection strategies. In addition to offering novel insights into microbial survival and dispersal, these findings have broader implications for biocontamination within the food, pharmaceutical, and medical sectors.

**Importance:** This study reveals that conidia / spores of a fungus Aspergillus calidoustus, which was isolated from spacecraft assembly cleanrooms, can survive simulated space-relevant stressors like ultraviolet irradiation, Martian cold atmospheric pressure, regolith exposure, ionizing radiation and specific doses of recommended dry heat sterilization for spacecrafts. Such fungal resilience demonstrates that the species can survive certain space and Mars conditions previously thought to be sterilizing, highlighting a need to revise current spacecraft decontamination standards that focus mainly on bacterial spores. This study also highlights the need for continued microbial monitoring of spacecrafts during transit from Earth to other planets not only to achieve goals of planetary protection but also to maintain healthy closed system for manned missions. Moreover, it is also alarming for an Earth origin fungal species due to biocontamination risks for food, medical, and pharmaceutical industries may require need for new standards of sterilization approaches transferable to space exploration.

## Introduction

Sterilization by radiation or heat exposure are critical protocols to a number of high-value industries, ranging from food safety (1, 2) and pharmaceuticals (3), to space sciences (4). However, some microorganisms exhibit remarkable adaptations, potentially allowing for survival under typically deleterious conditions. This has interesting ecological implications for many anthropological and terrestrial niches, as well as ecosystems beyond Earth. For instance, while interplanetary missions such as the active Mars rovers and the planned Mars Sample Return campaign (5) have the potential to yield some of the most exciting, contemporary scientific outputs, it must be ensured that they do not compromise the integrity of extraterrestrial ecosystems through the introduction of Earth-originating microorganisms (6).

Mars presents an array of hostile environmental factors that can significantly impact microbial survival; the thin Martian atmosphere is predominantly composed of carbon dioxide and provides only minimal protection from ultraviolet (UV) radiation, resulting in a surface-UV flux (100-280 nm) ∼10 times higher than on Earth (7). Additionally, the comparatively low surface pressure of ∼6 mbar (8), the lack of atmospheric water vapour, and the average annual Martian temperature of ∼ -60°C (9, 10), present a myriad of environmental challenges for any potential colonizing organism. However, various microorganisms are known to survive exposure to UV radiation (11–13), ionizing radiation (14–16), and the variable influence of desiccation and temperature extremes (17, 18). This suggests that there is potential for microbial survival in space or on Mars.

To mitigate these risks, the National Aeronautics and Space Administration (NASA) and other spacefaring agencies implement rigorous planetary protection standards, ensuring the cleanliness of extraterrestrial space and assembly cleanrooms (17, 19, 20). Germicidal techniques are typically quantified through enumerating aerobic spore-forming bacteria. *Bacillus pumilus* SAFR-032 is an example that has been extensively studied in the context of the space environment (11, 12). However, bacterial-spore detection alone may not be an accurate representation of bioburden (21). Fungi, such as *Aspergillus* and *Penicillium*, are common contaminants in spacecraft assembly facilities and have been isolated from Mars mission assembly cleanrooms (22). Fungal conidia also have the ability to endure extreme environmental stressors (23), including short-term exposure to limited simulated Martian conditions (19). It is therefore critically important to consider fungal conidia as a potential concern for planetary protection, and as valuable model species for understanding the adaptions of eukaryotic life to extreme environmental conditions.

This study embarks on a detailed assessment of microorganisms that could potentially serve as contaminants on Mars-bound spacecraft components. Unique fungal conidia, isolated from spacecraft assembly facilities (24), were compared against the radio-resistant microorganisms *Aspergillus fumigatus* (International Space Station strain (25), and *B. pumilus* (isolated from a spacecraft assembly facility (SAF) cleanroom (11). Microbial survival and morphology were assessed following exposure to simulated Martian conditions (SMC), in which a state-of-the-art experimental Martian simulation facility was used to expose microbial samples with a combination of the distinct Martian UV-radiation spectrum, atmospheric composition and pressure, and cooling to the mean Martian surface temperature. When exposed to UV radiation or extremes in temperature, microbial survival is influenced by the presence of soil (26, 27). As such, the influence of Martian regolith was also assessed during SMC experiments.

Prior to reaching Mars, contaminant organisms must survive two other potentially fatal events; an extended dose of ionizing space radiation accumulated during long-duration space travel (28), and exposure to spacecraft sterilization protocols. As such, microbial viability was also assessed following chronic exposure to a Californium-252 (^252^Cf) source, and after performing dry heat microbial reduction (DHMR; a common spacecraft-sterilization method (20). ^252^Cf is a radionuclide that provides a low-dose neutron field that closely approximates the chronic exposure patterns of galactic cosmic rays and solar particle events encountered inside spacecraft beyond low Earth orbit (28).

This research forms a comprehensive, end-to-end assessment of microbial survival during spacecraft sterilization, deep-space travel, and exposure to the Mars environment. This provides valuable insights into microbial resistance to different radiative environments and evaluates the impact of spacecraft sterilization. The survival of fungal species following high-energy UV exposure, extended ionizing-radiation doses, and sterilization protocols highlights the critical role of fungi in astrobiology and has broader implications sterilization techniques used in food science, the pharmaceutical industry and human health. This work is crucial in informing future planetary protection strategies and provides foundational knowledge into the tolerances of life to environmental extremes both on and beyond Earth.

## Materials and methods

### Isolate selection and sample preparation

A total of 29 microbial isolates were examined, including 27 fungal strains previously isolated from NASA Mars 2020 mission assembly facilities (24), as well as two spacecraft-associated organisms known to show high-levels of radiotolerance; *A. fumigatus* ISSFT-021-30 (25) and *B. pumilus* SAFR-032 (11). A complete list of isolates is included in Table 1 and Table S1.

**Table 1:** Microbial strains selected for exposure to SMC. . Isolation locations include NASA Jet Propulsion Laboratory – spacecraft assembly facility (JPL-SAF), Kennedy Space Center – payload hazardous servicing facility (KSC-PHSF), and the International Space Station (ISS). Survivability is recorded as positive (+) or negative (-) if, following SMC exposure, all replicates respectively demonstrated microbial growth or complete inactivation, respectively. Growth was detected in the sample preparation controls (dried sample, no irradiation) for all stains. *Aspergillus fumigatus* was not part of SAF but used as control.

| Species | Strain ID | Facility isolated | Survival after Martian irradiation (with/without regolith) |  |  |
| --- | --- | --- | --- | --- | --- |
|  |  |  | 5 min | 15 min | 30 min |
| *Aaosphaeria pasadenensis* | FJI-L9-BK-P1 | JPL-SAF (2018) | -/- | -/- | -/- |
| *Alternaria alternata* | FKII-L8-BK-P2B | KSC-PHSF (2018) | -/- | -/- | -/- |
| *Aspergillus calidoustus* | FKI-L3-BK-DRAB1 | KSC-PHSF (2018) | +/+ | +/+ | +/+ |
| *Aspergillus calidoustus* | FKII-L3-BK-PAB1 | KSC-PHSF (2018) | -/+ | +/- | +/+ |
| *Aspergillus* sp. | FKII-L3-BK-DR1 | KSC-PHSF (2018) | +/- | -/- | -/+ |
| *Aspergillus tubingensis* | FKII-L6-BK-DRAB1 | KSC-PHSF (2018) | +/- | -/- | -/+ |
| *Cladosporium crawfordii* | FJII-L10-SW-DR1 | JPL-SAF (2018) | -/- | -/- | -/- |
| *Cladosporium crawfordii* | FKII-L3-CM-PAB1 | KSC-PHSF (2018) | -/- | -/- | -/- |
| *Cladosporium endophyticum* | FKII-L2-CM-DR3 | KSC-PHSF (2018) | -/- | -/- | -/- |
| *Cladosporium prattii* | FJI-L7-BK-DG1 | JPL-SAF (2018) | -/- | -/- | -/- |
| *Neocatenulostroma microsporum* | FJI-L3-BK-P2 | JPL-SAF (2018) | -/- | -/- | -/- |
| *Penicillium cerradense* | FJI-L10-BK-P2 | JPL-SAF (2018) | -/- | -/- | -/- |
| *Talaromyces ruber* | FKI-L3-BK-DAB3 | KSC-PHSF (2018) | -/- | -/- | -/- |
| *Aspergillus fumigatus* | ISSFT-021-30 | ISS (2019) | +/+ | +/+ | +/+ |
| *Bacillus pumilus* | SAFR-032 | JPL-SAF (1999) | +/- | +/- | +/+ |

Purified conidia stocks were prepared by first plating each fungal isolate on Potato Dextrose Agar (PDA; Difco), and incubating at 25°C for a maximum of 8 weeks. Each plate was washed with 5 mL of sterile water (Molecular Biology Grade Water, Corning 46-000-CV), manually agitated for 1 min, and the resulting conidia suspension was harvested. Plate washing was repeated twice for each isolate and each suspension was allowed to settle for 3 h before the supernatant was discarded. Conidia were harvested via centrifugation (4500 G, 15 min, 25°C), and light microscopy was used to confirm the absence of fungal hyphae and the presence of >90% conidia. Similarly, a bacterial spore stock solution was prepared by culturing *B. pumilus* SAFR-032 (11) on Tryptic Soy Agar (TSA; Difco) at 25°C, manually transferring cultures to sterile Schaeffer Sporulation Medium (29), and continuing 25°C incubation for an additional 7 days. Spores were harvested via centrifugation (4500 G, 4°C, 15 min) and purified via triplicate washing (30). Light microscopy was used to ensure >99% refractile spores were present. Both the bacterial spore suspension and all purified conidia pellets were stored at 4°C.

Experimental samples were prepared by diluting fungal conidia or bacterial spores in sterile water to 10^7^ cells mL^-1^. 100 µL of each isolate suspension was deposited onto a precision-cleaned, spacecraft-grade, aluminium coupon (Al6061-T6 alloy, 13 mm diameter, 2 mm thickness) (31). Samples were allowed to dry under sterile conditions at 24°C, 32.5% relative humidity, for at least 12 h. To assess the effects of Martian regolith on fungal conidia or bacterial spore survival, a 1% (w/v) Martian regolith solution was prepared by suspending Martian regolith simulant (Atacama Desert soil) in sterile water, vortexing at high speed for 5 min, and collecting the supernatant after a 2 h separation period (32). The resulting Martian regolith solution was sterilized via autoclave. Fungal conidia or bacterial spores were then diluted to 10^7^ cells mL^-1^ in Martian regolith solution, before being deposited and dried on aluminium sample coupons in the same manner described above.

### Simulated Martian conditions

A preliminary UV-tolerance assay was performed, exposing dried isolate samples to a 3,000 J m^-2^ dose of UVC (CL-1000 UV crosslinker, UVP Inc, USA). Following this initial UV-tolerance screening, 13 unique SAF strains that demonstrated UVC tolerance (in addition to *A. fumigatus* and *B. pumilus*) were selected for analysis based on reported abundance on SAF cleanroom surfaces (22), a diverse selection of genera, and robust growth.

Selected strains were exposed to various SMC using experimental simulation facilities. Each strain was assessed with and without Martian regolith present, exposing samples to Martian irradiation for periods of 0, 5, 15 or 30 min. Simulated Martian UV radiation was provided with a collimated xenon arc lamp-equipped solar simulator (SciSun 300, Sciencetech, Canada), positioned 32 cm above the samples. The mean irradiance intensity was measured at the sample surface using calibrated miniature UV-Vis spectrometer equipped with a UV-enhanced optical fibre and cosine corrector (HR4000, QP115-2-XSR, CC-3-UV-S, Ocean Insight, Germany).

*Aspergillus calidoustus* FKI-L3-BK-DRAB1, *A. fumigatus* ISSFT-021-30 and *B. pumilus* SAFR-032 were selected for longer-term exposure to a broader range of Martian environmental conditions. In addition to assessing the influence of Martian regolith, samples were independently exposed to Martian atmospheric conditions (33) and up to 1440 min of Martian solar radiation. Prior to applying a simulated Martian atmosphere, the sample chamber was purged twice, removing ambient gas. A custom gas mixture, composed of 95.8% carbon dioxide, 2.1% argon, 1.9% nitrogen and 0.2% oxygen (Linde Gas, Germany), mixed using four mass flow control units (EL-FLOW Prestige, Bronkhorst, Netherlands), was then supplied at a constant flow rate of 6 mbar. The pressure was maintained using a needle valve and an adjustable vacuum pump (LVSF V, Welch, Germany), and monitored using a Pirani/cold cathode pressure gauge (PKR 360, Pfeiffer Vacuum, Netherlands). For samples not exposed to the simulated Martian atmosphere, a standard Earth atmospheric composition and pressure was maintained within the sample chamber.

Additional *A. calidoustus* samples were also exposed to the mean annual Martian surface temperature (–60°C) for up to 1440 min. This was achieved by cooling the copper sample plate with a liquid nitrogen flow-through system. To ensure the temperature reading accurately reflected the true temperature of the samples, a PT1000 temperature probe was adhered with thermally conductive epoxy (Loctite Stycast 2850FT, Henkel, Germany) to an identical aluminium coupon (13 mm diameter, 2 mm thickness) that the samples were deposited on. Thermal control was maintained using a Raspberry Pi (equipped with a MAX31865 RTD-to-digital converter) driving a proportional-integral-derivative (PID) feedback loop, which controlled the supply of liquid nitrogen via a solenoid valve. Samples not exposed to Martian temperatures were maintained at room temperature (21°C ±1°C).

### Neutron radiation exposure

To evaluate the impact of simulated space radiation on conidial survival rates, triplicate sample coupons of *A. calidoustus* were exposed to a chronic, low-dose ionizing radiation environment. The neutron radiation facility consisted of a sealed source panoramic irradiator (model 81-14R, J.L. Shepard, USA) containing an activity of 41.78 mCi ^252^Cf. The spontaneous fission of ^252^Cf provided a continuous source of fission neutrons, at low dose rates, which serve as a proxy for high-linear-energy-transfer radiation exposure, attributed to galactic cosmic rays (28). The exposure duration was varied according to the decay of the radiation source to provide a daily dose of 2.16 mGy over the trial period. Conidia sample coupons were exposed for a duration of 1 month and 6 months, and compared against control sample coupons (unirradiated samples stored for the duration of the exposure).

### Dry heat microbial reduction

DHMR was investigated as a means of sterilizing *A. calidoustus, A. fumigatus* and *B. pumilus* samples, based on previous methodologies (20, 34), with minor modifications. Briefly, triplicate sample coupons were aseptically loaded into sterile, blind-end, stainless-steel thermal spore exposure vessels (TSEV; 1.27 cm internal diameter, 10.16 cm length, 0.025 cm wall thickness), which were crimp-sealed with a silicone rubber septum (ST-495, Specialty Silicone Products, NY), held in place with 20 mm aluminium crimp-seals (Wheaton Science Products, NJ). Rubber septa were pierced with 22-gauge, side-hole needles (Hamilton Company, NV), and vacuum evacuated to 1.47 mbar (TriScroll 300 vacuum pump, Varian Vacuum Technologies, MA). TSEV were heated to either 125°C (a commonly used temperature in industrial sterilization processes (35) or 150°C (an approximate upper limit of thermal stress) using a high-temperature silicone oil bath (Model 6330, Hart Scientific, UT). After heating (∼90 s) samples were maintained at the target temperature for 5, 30, 60, 120, or 180 min.

After exposure, TSEV were placed in a room temperature water bath for 10 min. Sample coupons were aseptically removed from TSEV, and microbial survival was evaluated. In addition to the sample coupons, each DHMR run included a blank aluminium coupon as sterility control. Duplicate additional TSEV were inoculated but were not heated, serving as positive controls.

### Assessment of cell survival

Following the preliminary UV-tolerance screening, fungal sample coupons were submerged in 5 mL Potato Dextrose Broth (PDB; Difco) and incubated at 25°C for 7 days. Bacterial sample coupons were submerged in 5 mL Tryptic Soy Broth (TSB; Difco) and incubated at 25°C for 3 days. Following this incubation period, growth was qualitatively assessed based on turbidity.

Following SMC exposure, fungal conidia or bacterial spores were removed from sample coupons using polyvinyl alcohol extraction (36), and resuspended in 1 mL PBS. Quantification of viable microorganisms was then performed using either Most Probable Number (MPN) enumeration or plate counts (37), with serial dilutions in either PDB or TSB for fungal and bacterial samples, respectively. MPN assessment was performed in quadruplicate, incubating at 25°C observing turbidity after 72 h for bacteria and 1 week for fungi. Fungal conidia plate counts were performed by spreading 200 µL of each serial dilution to PDA, while bacterial plate counts employed 2 mL pour plating to TSA. Plates were incubated at 25°C for 7 days prior to enumeration of colony forming units (CFU).

Conidia and spore extraction of DHMR-or neutron radiation-exposed samples was performed as per previous methodologies (20). Briefly, sample coupons were submerged in 2 mL of a sterile water, with ∼150 mg spherical glass beads (1.27 mm diameter) then vortexed (Vortex-T, Genie 2) at maximum speed for 1 min. An additional 2 min sonication step (25 kHz, >0.35 W cm^-2^, Series 8500 Advanced Ultrasonic Generator, Branson Ultrasonics Corp., USA) was performed on bacterial samples. CFU were then enumerated via plate counting, as described above.

### Scanning electron microscopy imaging

Prior to performing scanning electron microscopy (SEM), dried samples on their aluminium coupon were Au/Pd sputter coated (Anatech Hummer 6.2, Anatech, USA). Argon gas was introduced to the sputter coating chamber at a pressure of <0.15 mbar, and 5 V was applied for 75s. Imaging was performed using an ultra-high-resolution Schottky SEM (SU7000, Hitachi High-Tech, USA) using conventional high vacuum conditions with a 1.00 kV beam and a working distance of 6.8-7.0 mm.

### Statistical analysis

MPN-based microbial enumeration results were analysed using MPNcalc (v. 1.5.0, Center for Food Safety and Applied Nutrition, USA). Parameters included six dilution steps, a 95% confidence level, and the asymptotic lognormal confidence interval technique.

For each SMC treatment, the mean log cell count of *A. calidoustus*, *A. fumigatus* and *B. pumilus* is plotted as a percentage, normalized against the respective control of each experiment (0 min Martian irradiation, Earth atmosphere, no Martian regolith, no thermal control). Results were compared using a one-way analysis of variance (ANOVA) with a Tukey Honest Significant Difference (HSD) post-hoc test (Table S2). The influence of Martian temperature is presented as a difference plot, displaying the change in mean cell count when compared to the non-thermally controlled equivalent. These results were assessed using an independent one-way ANOVA, comparing the normalized log cell survival percentage with the non-thermally controlled equivalent samples. The mean neutron radiation-induced reduction in viable cell number was compared for the 1- and 6-month exposure times, using a one-way ANOVA. Cell survival following 125°C DHMR was plotted as the log cell number over sterilization time. Plots were constructed and statistical analyses were performed using RStudio (v. 2023. 12.0+369, Postit Software, USA).

## Results

### Strain selection

Among 29 fungal isolates examined 25 strains survived exposure to 3000 J m^-2^ of UVC. All UVC-resistant fungi belonged to filamentous ascomycete fungi. During the down-selection process based on UV resistance and species diversity, 13 fungal strains representing 12 species across seven genera were selected (Table 1). Additionally, the UV-resistant bacterium *Bacillus pumilus* SAFR-032 and a fungus isolated from the International Space Station (ISS) were included as control for further exposure in the experimental Mars simulation chamber. Isolates that were excluded from further analysis are documented in Table S1.

### Mars simulation chamber operating principles

The experimental Mars simulation facility is a compact and cost-effective means for the independent replication of Martian environmental parameters including solar irradiation, atmospheric parameters and cooling to surface temperatures. The housing consists of a gas-tight, ∼900 cm^3^ aluminium chamber fitted with a 6.3 cm diameter MgF_2_ viewport (Fig. 1a), which is highly transmissive to wavelengths as low as 120 nm. The solar simulator can provide UV-illumination corresponding to Martian locations ranging from the equator to the polar regions, or from the planetary surface to the upper atmosphere. In this experiment, samples were irradiated with a UV spectrum comparable to the solar radiation present at the Martian surface at 0° N (Fig. 1b), which when integrated from 200-400 nm, provided a mean intensity of 56.01 W m^-2^ (±3.75 W m^-2^). This corresponds well with previous Mars solar irradiance studies (7, 19).

**Fig. 1:**
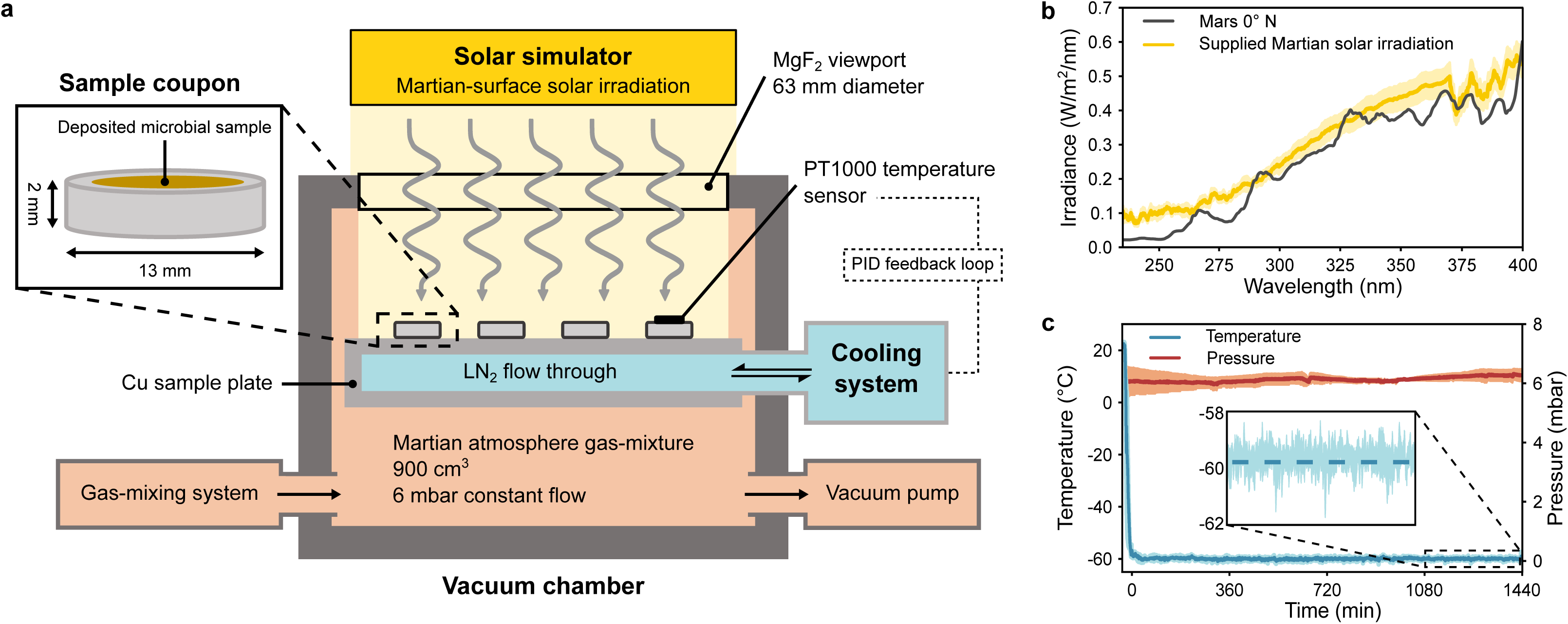
Configuration of the experimental Mars simulation chamber. **a**. Schematic representation of the experimental facilities used to expose samples deposited on aluminium coupons (inset) to SMC. **b**. Mean spectrum of supplied Martian solar irradiation (yellow, n=6), compared against calculated Martian surface solar irradiation at 0°N [7] (black). **c**. Mean temperature (blue, n=2) and pressure (orange, n=2) measured during a 1440 min Mars simulation period. Inset shows fine-scale temperature variations within the final 360 min period. Shaded regions show standard deviations.

The vacuum and gas-supply capabilities of the simulation chamber allows for experiments to be conducted with a constant supply of any user-defined gas mixture, or under vacuum conditions (as low as 5×10^-2^ mbar). This experiment utilized a customized Martian atmosphere gas-mixture, supplied at a constant flow rate maintaining a pressure of 6 mbar (±0.03 mbar; Fig. 1c). Active liquid-nitrogen cooling and passive heating to room temperature allows for thermal control via the copper sample plate (Fig. 1a). The internal temperature probe - PID feedback loop can be used to execute customize thermal cycles (i.e., Martian day-night cycles) or to maintain a static temperature. This work maintained a mean sample temperature of -59.4°C (±0.6°C) over the course of 1440 min exposures. This temperature was achieved following a mean cooling time of 18 min (Fig. 1c).

### Exposure to simulated Mars conditions

All tested strains of *Aspergillus* (as well as *B. pumilus*) had the capacity to survive up to 30 min of Martian solar radiation exposure, however, the presence of Martian regolith did not appear to have a consistent influence on survival. In contrast, the minimum dose (5 min) of simulated Martian solar radiation was lethal to all other examined fungal strains, regardless of the presence of Martian regolith (Table 1). Two strains of *Aspergillus* that demonstrated consistent survival, *A. calidoustus* FKI-L3-BK-DRAB1 and *A. fumigatus* ISSFT-021-30, along with *B. pumilus* SAFR-032, were selected for quantitative analysis following exposure to a broader set of Martian environmental variables.

*A. calidoutus* exhibited a particularly strong tolerance to Martian solar UV, surviving a 1440 min dose, with a 3-log mean reduction in cell number across all treatments (Fig. 2a). Exposure to a simulated Mars atmosphere negatively influenced the survival of *A. calidoutus*, with this trend being most prominent under lower irradiation times. Exposure to the Martian atmosphere without irradiation resulted in a 4.5 times reduction in cell survival, however the presence of regolith buffered this effect to a 1.6 times cell survival reduction (Fig. 2a). Other than this instance, Martian regolith did not have a clear influence on the growth of *A. calidoutus*.

**Fig. 2:**
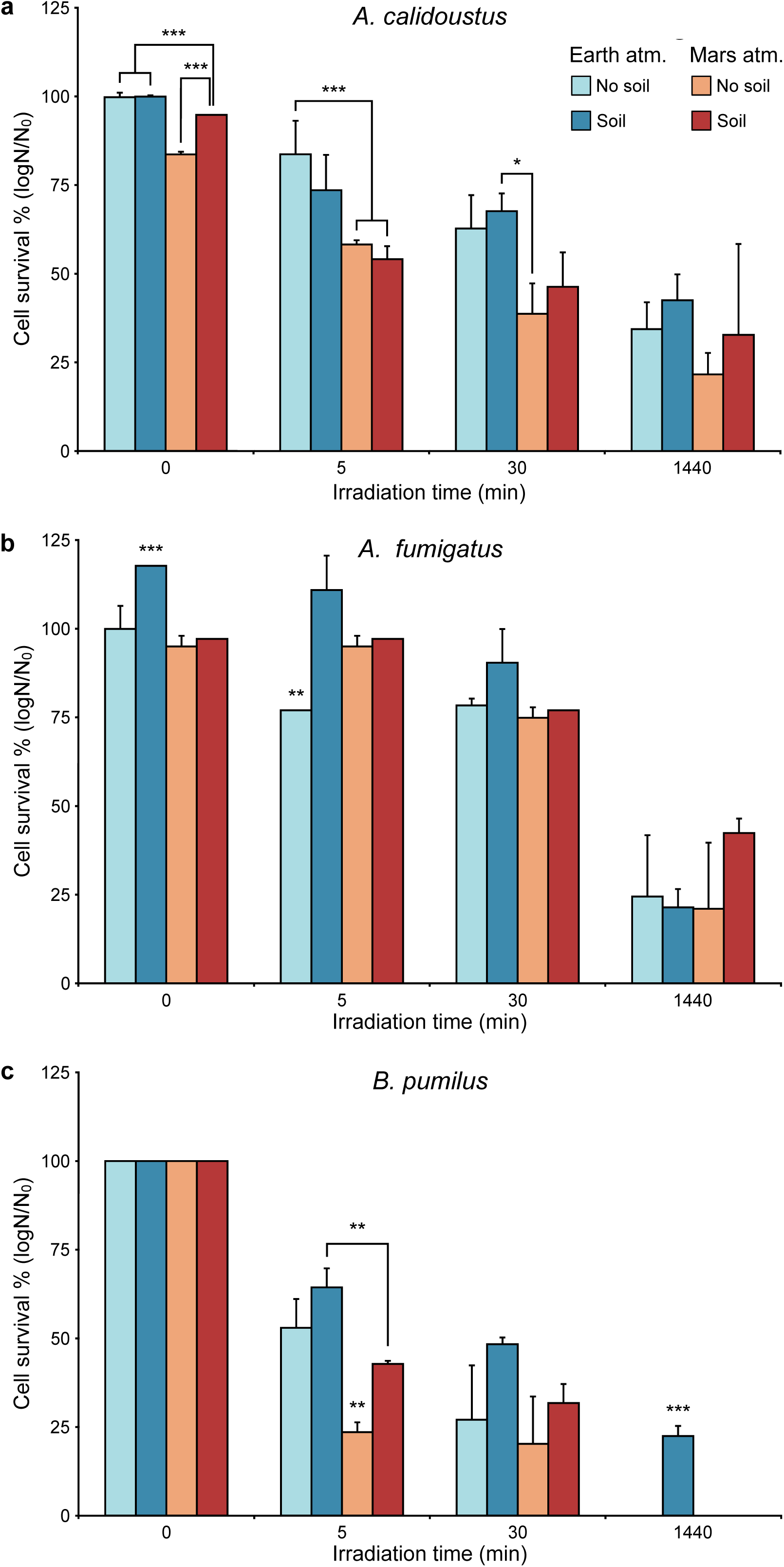
Mean cell survival following exposure to simulated Martian conditions. Microbial samples were irradiated with a simulated Martian solar spectrum whilst simultaneously being exposed to either an Earth atmosphere (blue) or Martian atmospheric conditions (red). Samples were comprised of either pure dried microbial cells (light shade) or a mix of dried cells and Martian regolith (dark shade). **a.** *A. calidoustus* conidia. **b.** *A. fumigatus* conidia. **c.** *B. pumilus* spores. n=3 and error bars represent standard deviation. Asterisks indicate treatments that statistically differ, as determined by a one-way ANOVA (p-value <0.05 = *, <0.01 = **, <0.001 = ***).

*A. fumigatus* had a similarly high tolerance to Martian solar UV exposure. Across all treatments a 1440 min irradiation dose resulted in a ∼3 order of magnitude reduction in cell number. However, *A. fumigatus* was more resilient to lower-dose irradiations than *A. calidoustus*, with ∼5×10^3^ cells still being present after both 5- and 30-min Martian solar irradiations (Fig. 2b). The presence of regolith significantly enhanced the survival of *A. fumigatus* under an Earth atmosphere. In contrast, the survival of *A. fumigatus* cells exposed to Mars-atmosphere conditions was unaffected by the presence of Martian regolith (Fig. 2b).

Simulated Martian irradiation exposure was more detrimental to *B. pumilus* spores than either *Aspergillus* species, and both the Mars atmosphere and regolith prominently influenced the survival of irradiated *B. pumilus* (Fig. 2c). Specifically, Martian regolith promoted survival, increasing cell numbers by ∼1-log across both the 5- and 30-min irradiation doses. In contrast, the Mars atmosphere increased the radiation-susceptibility of *B. pumilus*. These trends are present across all irradiation treatments, with the combined influence of Martian regolith and atmosphere being best exemplified after the 1440 min irradiation exposure; this dose was lethal for all *B. pumilus* samples, except for those both combined with regolith and under an Earth atmosphere (Fig. 2c).

During a 1440 min exposure of *A. calidoustus* to a broader suit of SMC, the combined influence of cooling to -60°C (the mean Mars surface temperature), Martian solar UV radiation and the Mars atmosphere resulted in a complete loss of cell viability, with Martian regolith providing no protective capacity (Fig. 3a). Without irradiation, cell survival was observed, but was highly influenced by the combined influence of temperature, atmosphere and regolith. Under an Earth atmosphere, cold exposure induced a ∼30-fold decrease in viability. When exposed to the Martian atmosphere however, the deleterious effect of cold exposure was ∼10 times less severe. When Mars-atmosphere exposed samples were compared against their non-thermally controlled equivalents, those lacking Martian regolith demonstrated 3 times higher survival when exposed to Mars surface temperatures.

**Fig. 3:**
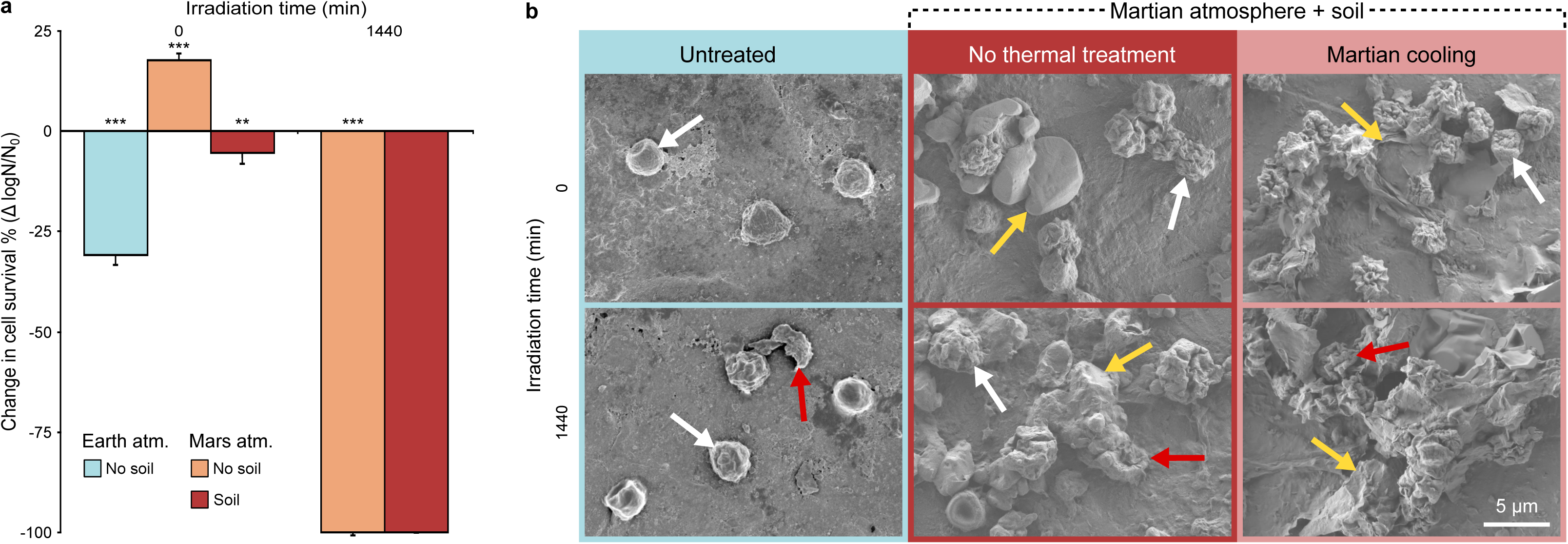
Influence of simulated Martian temperature on the survival and morphology of *A. calidoustus* conidia. **a.** Change in mean survival induced by 1440 min of Martian cooling in pure dried microbial cell samples (light red) or dried cells mixed with Martian regolith (dark red). Simultaneous to Martian cooling, samples were exposed to up to 1440 min of simulated Martian solar irradiation and Martian atmosphere. This was compared against unirradiated pure-cell samples, under an Earth atmosphere (light blue). n=3 and error bars represent standard deviation. Asterisks indicate treatments that statistically differ from their non-thermally treated equivalent, as determined by a one-way ANOVA (p-value <0.05 = *, <0.01 = **, <0.001 = ***). **b.** SEM images demonstrating the morphological changes of *A. calidoustus* conidia during exposure to SMC (irradiation, atmosphere, regolith and cooling). White arrows indicate intact conidia, red arrows indicate lysed conidia, and yellow arrows indicate soil particles. Scale bar equals 5 µm.

**Fig. 4:**
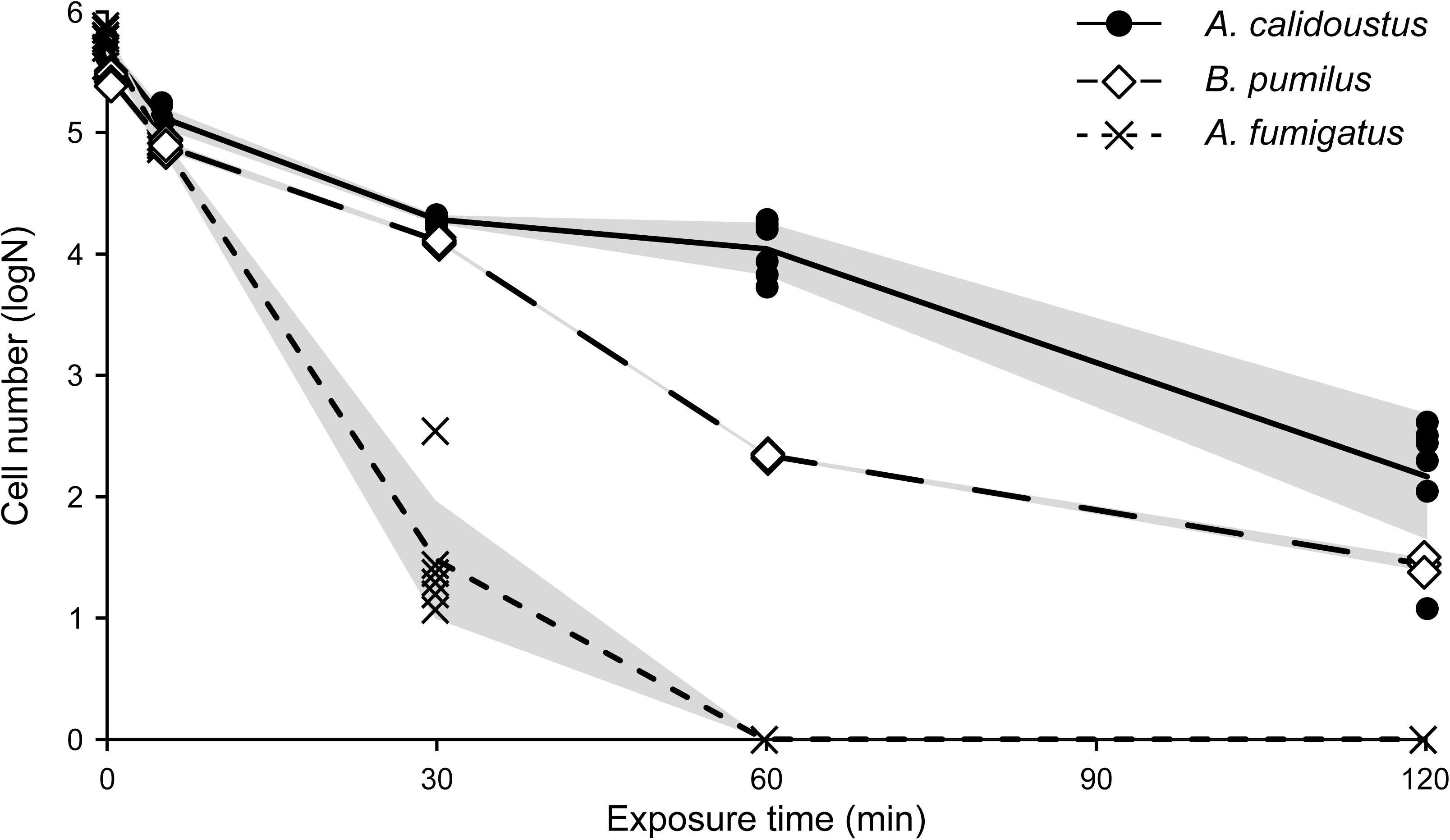
Survival following 125°C DHMR. Individual data points are shown while lines represent the mean cell number (black circle with solid line = *A. calidoustus*, white diamonds with heavy dashed line = *B. pumilus*, black crosses with fine dashed line = *A. fumigatus*). n=6 and shaded regions show standard deviations.

SMC induced significant morphological changes on *A. calidoustus* conidia. Prior to exposure, the majority of conidia exhibited a smooth surface with occasional cells having a concave dent present. Following a 1440 min irradiation period, a large proportion of conidia sporadically ruptured, with almost all intact conidia showing some surface irregularities (Fig. 3b). In the presence of Martian regolith, conidia adhered to soil particles and readily formed clusters. *A. calidoustus* exposed to the Martian atmosphere frequently showed deep pitting or scaring of the conidia surface. The combined influence of the Martian atmosphere and irradiation resulted in a combination of conidia deformation and extensive ruptures (Fig. 3b). Cooling to -60°C did not appear to further alter the morphology of *A. calidoustus*, however, the previously smooth Martian regolith particles took on a rough, sheet-like appearance. Very few intact conidia were observed following exposure to the full suite of SMC, with the remaining conidia being particularly difficult to distinguish from the surrounding regolith (Fig. 3b).

### Neutron radiation exposure

Samples of *A. calidoustus* exhibited high resilience to long-duration neutron-radiation exposure, however, a dose-dependent decline in viable conidia did occur (F_(1, 4)_ = 1232.7, p < 0.001). Specifically, 1-month neutron-radiation exposure decreased cell counts by 4.5 x10^5^, whereas the 6-month exposure period resulted in a loss of 10.3 x10^5^ viable cells. Respectively, these equate to 35% and 57% viable conidia reductions, when compared against non-exposed controls.

### Dry heat microbial reduction

When exposed to a 125°C DHMR regime, *A. calidoustus* exhibited significantly higher resilience than either *A. fumigatus* or *B. pumilus*. Although, the survival of *A. calidoustus* declined logarithmically and the 120 min DHMR period resulted in a 3-log loss in viability, compared to positive controls. During 5- and 30-min exposures of 125°C DHMR *B. pumilus* demonstrated a similar response as *A. calidoustus*, however a greater reduction (>3-logs) was observed by 60 min of exposure. After 120 min, *B. pumilus* spores were still present, but the viable number had reduced by 4-log. In contrast, *A. fumigatus* was more susceptible to DHMR at 125°C, with complete inactivation of *A. fumigatus* occurring between 30 and 60 min of exposure. In contrast, the 150°C DHMR exposure was lethal to all three isolates after only a 5 min period. Negative controls remained free from growth.

## Discussion

This study provides a unique and comprehensive assessment of microbial resilience, investigating the impact of multiple extreme stressors, from spacecraft bioburden reduction to the experimental simulation of space radiation and Martian environmental conditions. 23 SAF-isolated fungal strains were capable of surviving UVC exposure, of which several strains of *Aspergillus* also demonstrated tolerance to further doses of simulated Martian irradiation. Specifically, *A. calidoustus* (FKI-L3-BK-DRAB1) exhibited exceptional resistance to an extended dose of Martian irradiation, demonstrated comparable survival as the radiotolerant fungi *A. fumigatus*, and exceeded the survival of *B. pumilus* spores, which have been proposed as a UV-resistant bioindicator species [28]. Complete deactivation of *A. calidustous* conidia was only achieved through combined, prolonged exposure to both Martian irradiation and cooling.

Exposure to SMC revealed a variable and complex interplay between these environmental factors. The high concentration of CO_2_ and low pressure of the Martian atmosphere did not clearly influence *A. fumigatus*, while both *A. calidoustus* and *B. pumilus* demonstrated a general trend of decreased survival under the Mars atmosphere. In the absence of Martian irradiation, cooling under a Mars atmosphere increased the survival of *A. calidoustus*. However, the potential cryoprotective effect of the CO₂-rich Martian atmosphere was negated by the presence of the Martian regolith (possibly due to increased thermal conductivity enhancing microbial stress). In contrast, the presence of Martian regolith had minimal to no effect on the survival of fungal conidia exposed to Martian irradiation. As radiative shielding from Martian soil is in-part dictated by particle size (26), it is possible that the Martian regolith solution used in this study (soil particles <10 µm, see Fig. 3b) fell below the soil particle size-threshold capable of providing radiative protection. The combined influence of different simulated Martian environmental factors on survival, was also correlated with changes in cellular morphology. These result are inline with previous studies that have also demonstrated a variable, species- or strain-specific influence of elevated CO_2_ concentrations, cold temperatures and humidity (38, 39), and emphasize the complex relationship between the various Martian environmental factors and how they influence microbial survival and morphology.

In addition to surviving a broad suit of SMC, *A. calidoustus* conidia also exhibited >40% viability following a 6-month neutron radiation exposure. The ²⁵²Cf neutron source used in this experiment resulted in a chronic radiation dose of 2.16 mGy day^-1^, closely matching modelled interplanetary dose rates (∼390 mGy total over several months) (28). Prior simulated space radiation studies have typically employed acute, high-dose exposures, often within minutes. For example, the extremotolerance of *Aspergillus niger* spores was demonstrated following exposure to X-rays (LD₉₀ = 366 Gy), helium ions (LD₉₀ = 506 Gy), and iron ions (LD₉₀ = 112 Gy) (40). However, such findings may be understating fungal tolerance to sustained, low dose-rate exposures, which more accurately reflect spaceflight conditions. As such, we recommend future studies of this nature to adopt a time-resolved approach, examining extremotolerant organisms under chronic exposure conditions that mirror real mission scenarios.

DHMR is an important sterilization process, particularly useful for material surfaces that would be degraded by UV light, or are in shaded areas (41). Following 125°C DHMR, *A. calidoustus* demonstrated equivalent early-stage resistance and exceeded the late-stage resistance of *B. pumilus* spores. In contrast, *A. fumigatus* was completely inactivated within 30 min of 125°C DHMR exposure. To completely eradicate *A. calidoustus* conidia, 150°C DHMR exposure was required. NASA’s current dry heat sterilization guidelines range from 110°C to 126°C (42), and traditionally planetary protection protocols primarily focus on bacterial spores. As such, this work highlights the need for alternative spacecraft-sterilization methods and a greater consideration of fungal species as bioburdens.

Fungal species of this nature also pose risks in contexts beyond space science. In addition to *A. calidoustus* demonstrating resistance to the variety of environmental conditions and common sterilization techniques explored in this study, many other members of the *Aspergillus* genera can remain viable following pasteurization or heat-treatment (43); techniques often relied upon in the pharmaceutical industry or food-safety sector. Furthermore, *Aspergillus* is known to cause number of health complications, often associated with respiratory infections including COVID19 (44) and COPD (45). As such, understanding the tolerances of fungal conidia is widely important, with the results of this work having implication across a range of industries.

This is the first study to perform an end-to-end evaluation of multiple spaceflight associated environmental stressors on SAF-isolated microbial species. This work builds upon the studies that have demonstrated experimental planetary- and space-simulation facilities as essential tools for future astrobiology studies [31, 32]. As *A. calidoustus* (FKI-L3-BK-DRAB1) evaded standard spacecraft sterilization, survived an equivalent ionizing radiation dose to that accumulated during a six-month journey to Mars, and remained viable following SMC exposure (with only the combination of Martian cooling and irradiation being lethal), this strain is one of the most likely candidates for forward contamination. Fungal conidia are often overlooked despite exhibiting comparable, if not superior, resilience than bacterial spores. Their presence in cleanroom environments, in combination with the results of this study underscore fungal conidia as a significant consideration for planetary protection. We support the concept that aerobic bacterial spore detection alone is not sufficient for cleanroom bioburden monitoring, and that a methodological paradigm shift is required in the aeronautical, pharmaceutical and medical industries (46). Fungal conidia and their potential interaction with localized Martian microenvironments must be considered in future microbial-contamination mitigation protocols. To continue providing valuable insights into the limits of eukaryotic life and will contributing to the broader understanding of microbial adaptability, future astrobiology research must consider both the genetic and physiological responses of fungi under extreme environments.

## Acknowledgements

Part of the research described in this publication was carried out at the Jet Propulsion Laboratory, California Institute of Technology, under a contract with the National Aeronautics and Space Administration (80NM0018D0004). We acknowledge NASA grants 80NSSC23K1298 and NNX15AK13G which cover the neutron facility at Colorado State University at Fort Collins. We would like to thank Snehit Mhatre for isolating the strains. We thank Biotechnology and Planetary Protection Group members, especially Cynthia Ly for collecting samples.

## Author contributions

KV, AE, WS and NKS conceptualized the project, with inputs on experimental design from AMC and DJB. AMC and GF performed sample preparation and processing for post-exposure cell enumeration for SMC experiments. SW and RN constructed the Martian simulation chamber. DJB, with the assistance of SW and RN, performed SMC sample exposure and *in-situ* measurements. JJB, AB and MW performed neutron radiation exposure experiments. WS performed neutron-radiation exposures. DJB and AMC performed data analysis and wrote the original manuscript, which was edited by KV. All authors contributed to, and approved the final manuscript.

## Data availability

The data underlying this article are available in the manuscript and in its supplementary material. Novel species have previously been deposited in the German Collection of Microorganisms in Cell Cultures (DSM 114620, 114621, 114623, 114624, 114626, 114627, 114728, 114631) or the Agricultural Research Service Culture Collection (NRRL 64422, 64423, 64424, 64425, 64427, 64428, 64432, 64433, 64434).

## Informed consent statement

This manuscript was prepared as an account of work sponsored by NASA, an agency of the US Government. The US Government, NASA, California Institute of Technology, Jet Propulsion Laboratory, and their employees make no warranty, expressed or implied, or assume any liability or responsibility for the accuracy, completeness, or usefulness of information, apparatus, product, or process disclosed in this manuscript, or represents that its use would not infringe upon privately held rights. The use of, and references to any commercial product, process, or service does not necessarily constitute or imply endorsement, recommendation, or favouring by the US Government, NASA, California Institute of Technology, or Jet Propulsion Laboratory. Views and opinions presented herein by the authors of this manuscript do not necessarily reflect those of the US Government, NASA, California Institute of Technology, or Jet Propulsion Laboratory, and shall not be used for advertisements or product endorsements. © 2025. All rights reserved

---

**Table S1.**
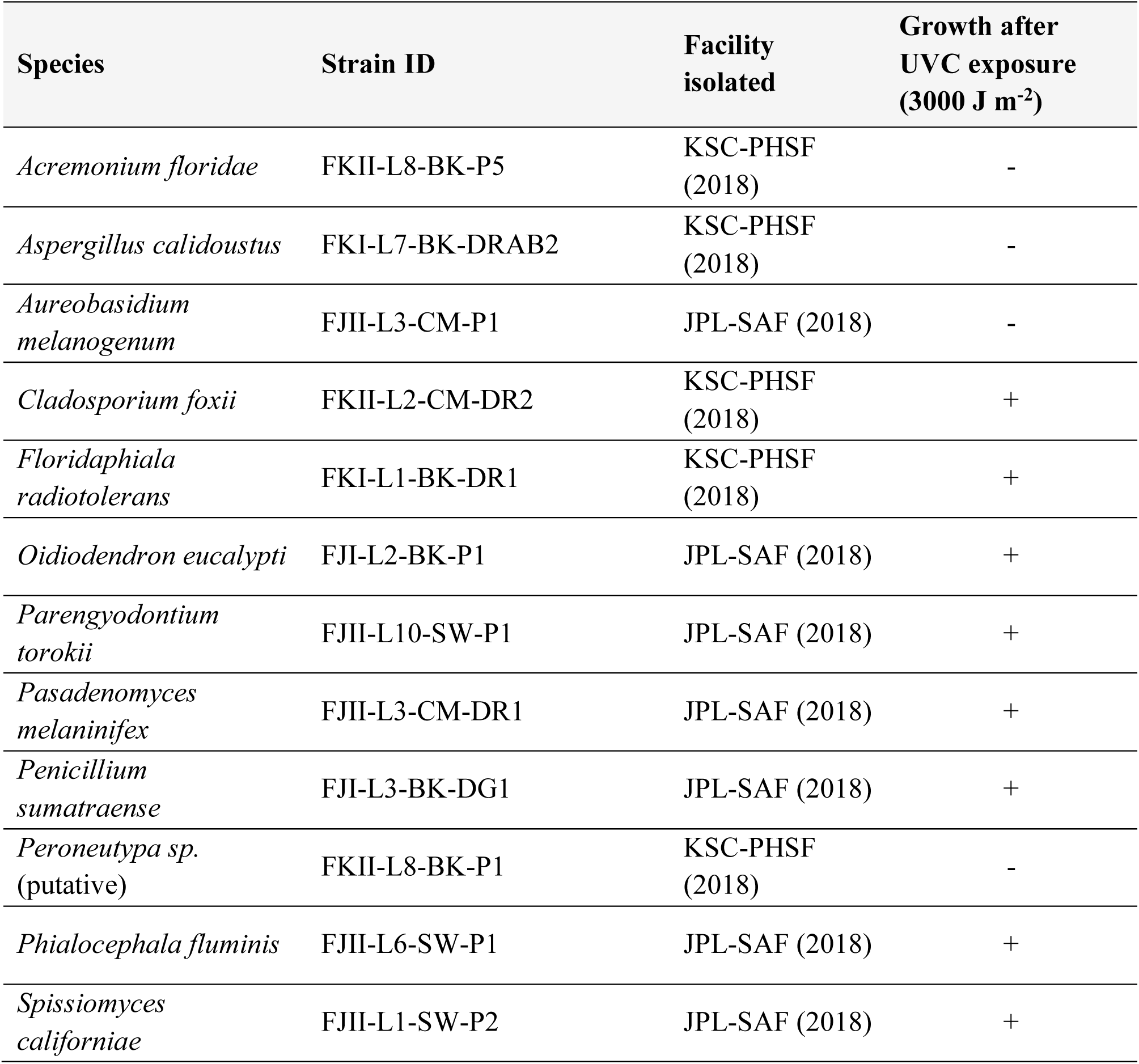
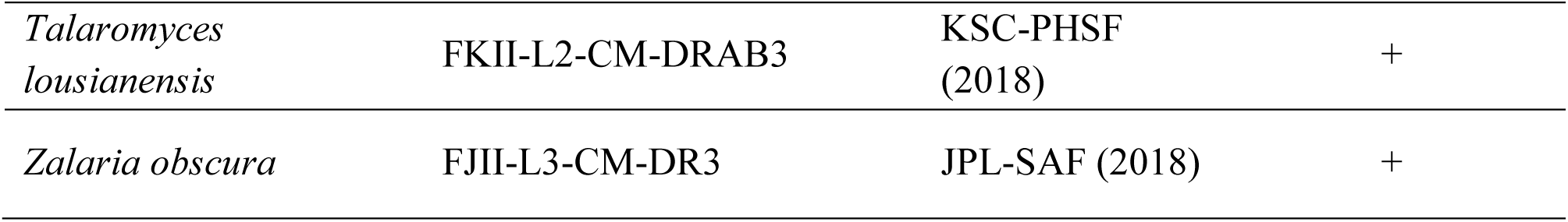
Microbial isolates excluded from SMC exposure. Isolation locations include Kennedy Space Center – payload hazardous servicing facility (KSC-PHSF) and NASA Jet Propulsion Laboratory – spacecraft assembly facility (JPL-SAF). Survivability is recorded as positive (+) or negative (-) if all replicates respectively demonstrated microbial growth or complete inactivation, respectively.

**Table S2.**
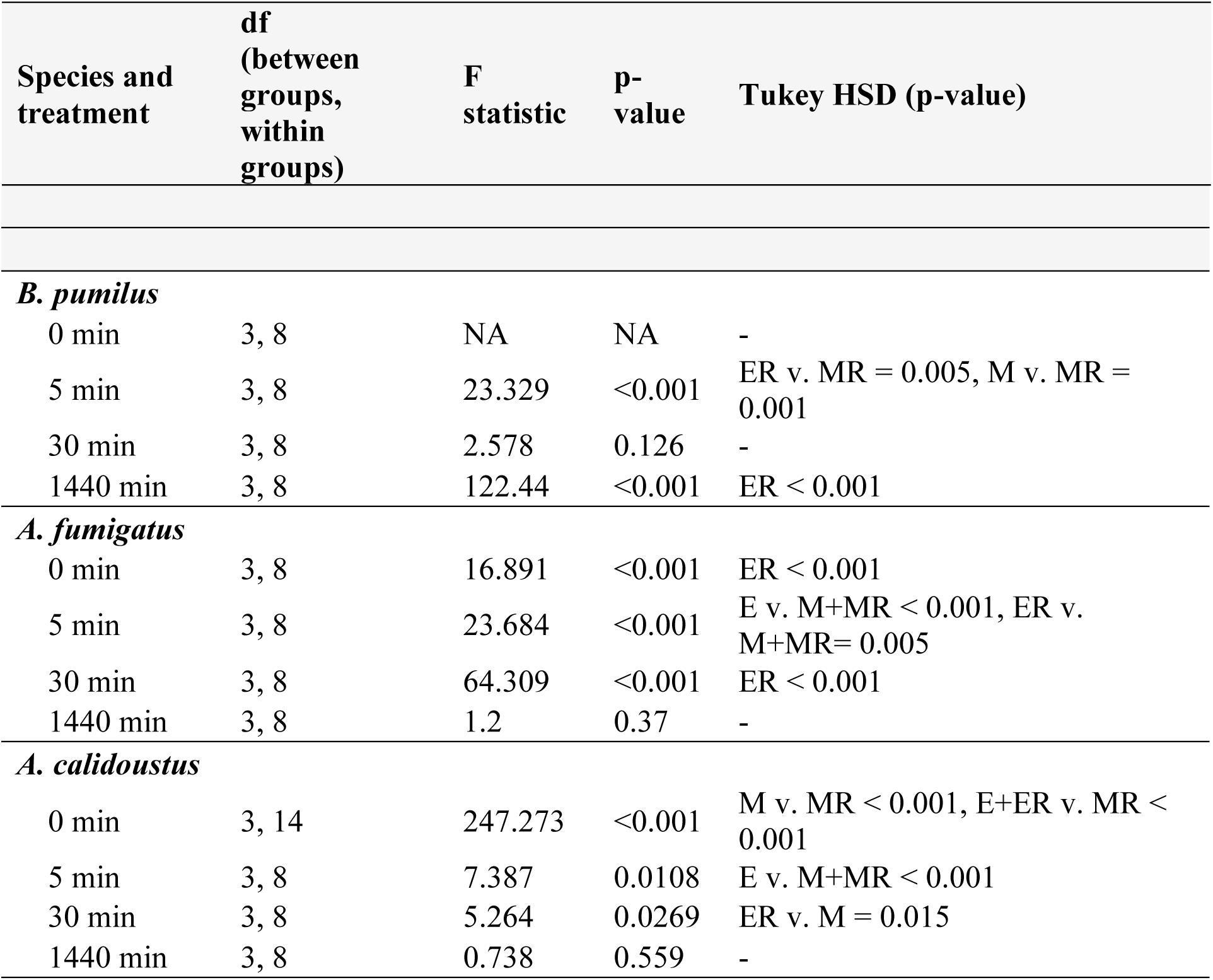
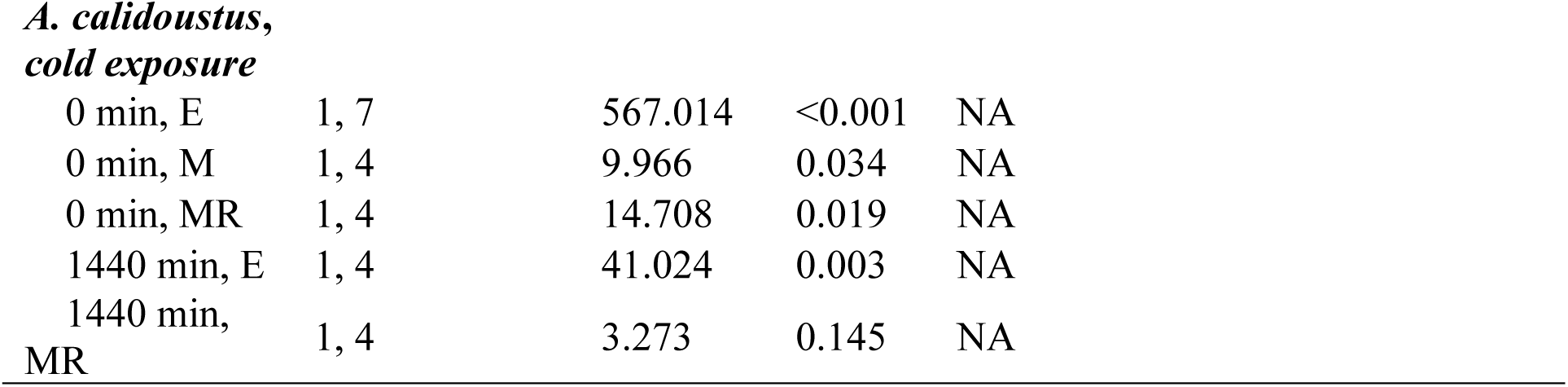
One-way ANOVA results. Statistical comparisons were made between each species’ irradiation exposure treatment group for. For cold-exposed *A. calidoustus* samples, comparisons were made between against each respective non-temperature controlled treatment. Environmental treatments are indicated using the following abbreviations: E = Earth atmosphere, M = Martian atmosphere, R = Martian regolith.

